# Exercise is associated with younger methylome and transcriptome profiles in human skeletal muscle

**DOI:** 10.1101/2022.12.27.522062

**Authors:** Sarah Voisin, Kirsten Seale, Macsue Jacques, Shanie Landen, Nicholas R Harvey, Larisa M Haupt, Lyn R Griffiths, Kevin J Ashton, Vernon G Coffey, Jamie-Lee M Thompson, Thomas M Doering, Malene E Lindholm, Colum Walsh, Gareth Davison, Rachelle Irwin, Catherine McBride, Ola Hansson, Olof Asplund, Aino E Heikkinen, Päivi Piirilä, Kirsi H Pietiläinen, Miina Ollikainen, Sara Blocquiaux, Martine Thomis, Dawn K Coletta, Adam P Sharples, Nir Eynon

**Author notes:** Corresponding Authors: Dr Sarah Voisin, Institute for Health and Sport (iHeS), Victoria University, Melbourne, Australia 8001; Professor Nir Eynon, Institute for Health and Sport (iHeS), Victoria University, Melbourne, Australia 8001.

## Abstract

Exercise training prevents age-related decline in muscle function. Targeting epigenetic aging is a promising actionable mechanism and late-life exercise mitigates epigenetic aging in rodent muscle. Whether exercise training can decelerate, or reverse epigenetic aging in humans is unknown. Here, we performed a powerful meta-analysis of the methylome and transcriptome of an unprecedented number of human skeletal muscle samples (n = 3,176). We show that: 1) individuals with higher baseline aerobic fitness have younger epigenetic and transcriptomic profiles, 2) exercise training leads to significant shifts of epigenetic and transcriptomic patterns towards a younger profile, and 3) muscle disuse “ages” the transcriptome. Higher fitness levels were associated with attenuated differential methylation and transcription during aging. Furthermore, both epigenetic and transcriptomic profiles shifted towards a younger state after exercise training interventions, while the transcriptome shifted towards an older state after forced muscle disuse. We demonstrate that exercise training targets many of the age-related transcripts and DNA methylation loci to maintain younger methylome and transcriptome profiles, specifically in genes related to muscle structure, metabolism and mitochondrial function. Our comprehensive analysis will inform future studies aiming to identify the best combination of therapeutics and exercise regimes to optimize longevity.

## Introduction

The United Nations has declared 2021-2030 the “Decade of Healthy Ageing” to assist the aging population in living healthier for longer. Identifying reliable aging biomarkers that can be targeted by longevity-promoting interventions is a global priority, however, requires sizable human cohorts across a broad range of ages and relevant tissues, which is both costly and time-consuming. The last two decades have seen an open science revolution with the creation of free-access repositories overflowing with molecular data from all levels of gene regulation (e.g. epigenomics, transcriptomics). These rich repositories allow for the exploration of aging mechanisms and their susceptibility to environmental stressors, even in healthy and/or young individuals, since age-related changes gradually accumulate from early life and affect organ systems years before disease manifestation^1^.

Aging is associated with a loss of muscle mass and function that leads to increased adverse outcomes including falling injury, functional decline, frailty, earlier morbidity and mortality^2^. Exercise training is one of the most affordable and effective ways to promote healthy aging^3^, as being physically active reduces mortality from all causes, independent of levels and changes in several established risk factors (overall diet quality, body mass index, medical history, blood pressure, triglycerides, and cholesterol)^4^. Cardiorespiratory fitness, as estimated by maximal oxygen uptake (VO_2max_) during an exercise test, shows a strong, graded, and inverse association with overall mortality^5^. However, we have an incomplete understanding of the fundamental mechanisms by which physical activity delays the age-related decline in skeletal muscle function.

At the molecular level, aging arises from a tip in the balance between cellular damage and compensatory mechanisms^6,7^. Cells undergo constant damage, such as genomic instability, telomere attrition, epigenetic alteration, loss of proteostasis, and disabled macroautophagy^6^. This leads to impaired nutrient sensing, mitochondrial dysfunction and cellular senescence, which play more nuanced roles in the aging process, as they can be beneficial at a young age (e.g. the nutrient-sensing network contributes to organ development until young adulthood but can have a detrimental role beyond this stage). Low doses such as occurs in mitochondrial dysfunction can stimulate beneficial counterreactions via mitohormesis, or if spatially confined (e.g. cellular senescence suppression of oncogenesis and improved wound healing)^6^. Eventually, the accumulated damage inflicted by these primary and antagonistic hallmarks can no longer be compensated, leading to stem cell exhaustion, altered intercellular communication, chronic inflammation and dysbiosis, which are ultimately responsible for the physiological decline associated with aging^6^. The effect of aging on DNA methylation (DNAm) patterns are so profound that machine learning has spawned highly accurate predictors of both chronological and biological age (termed “epigenetic clocks”)^8^. We developed an epigenetic clock for human skeletal muscle^9^ and reported widespread changes in the muscle methylome at genes involved in muscle structure and function^10^. Downstream of epigenetic processes, changes in transcriptional patterns at genes involved in central metabolic pathways and mitochondrial function have also been reported in muscle during aging^11^. Regular exercise mitigates the age-related loss of proteostasis^12,13^, mitochondrial dysfunction^3,14^, and stem cell exhaustion^3,15^ in muscle, but there is currently limited evidence for its effect on age-related epigenetic and transcriptomic changes.

Several cross-sectional analyses found only weak associations between self-reported physical activity levels and epigenetic age in blood^16–18^ or skeletal muscle^18^, after adjusting for confounders such as diet. However, self-reported measures of physical activity poorly reflect actual physical activity levels, particularly in individuals with higher body fat and females who typically overestimate energy expenditure^19^. Furthermore, these studies relied solely on DNAm clocks to quantify age-related changes in epigenetic patterns. While aging is associated with widespread changes at a plethora of CpG sites, epigenetic age, as measured by epigenetic clocks, is a single value that encompasses a very small portion of the aging methylome (typically a few hundred age-related DNAm loci, also called CpGs). Therefore, it offers a very narrow and incomplete view of the aging methylome, and exercise training may affect aging regions that are not captured by epigenetic clocks. Interestingly, a recent study found evidence that late-life exercise mitigates age-related epigenetic changes in mouse gastrocnemius muscle^20^. In this study, the authors did not use clocks but investigated all DNAm loci that change with age in mouse muscle and applied a direct exercise training intervention. A couple of human studies are in line with these results: resistance training was shown to offset age-related changes both in the nuclear^21,22^ and mitochondrial^23^ epigenome. Unfortunately, these studies had small sample sizes and have not been replicated to date, so they should be regarded as preliminary^24^. They were also restricted to resistance training in males, which does not speak for the effect of exercise training in general in males or females across the lifespan. In another, large-scale meta-analysis, age-related changes in the transcriptome showed an inverse correlation with higher cardiorespiratory fitness (CRF)^11^. However, these associations were cross-sectional and could be confounded by other environmental and lifestyle factors that co-occur with higher CRF levels (e.g. a better diet). Therefore, there is a great need for a comprehensive, integrative and robust assessment of the effects of CRF, exercise training and inactivity on age-related molecular changes in human muscle.

To address these limitations, we identified relevant published datasets from online databases that we combined with our own original data to characterise the effect of cardiorespiratory (CRF), exercise training and inactivity on human skeletal muscle aging across the methylome and transcriptome. We first compiled a list of age-related changes in 1,251 samples across 16 cohorts (DNAm) and 1,925 samples across 21 cohorts (mRNA expression). We then tested cross-sectional associations between aging molecular profiles and CRF; we hypothesized that individuals with higher CRF would display ‘younger’ profiles than expected at age-related CpGs and mRNAs. Then, we investigated directly whether exercise training could shift molecular profiles in human skeletal muscle towards a younger profile, using high-resolution longitudinal data collected from exercise training studies of various types and durations. Finally, we tested whether inactivity could ‘age’ transcriptomic profiles in human skeletal muscle, using longitudinal data from forced immobilization interventions in humans. This work provides a comprehensive and integrative map of the effect of physical activity on the age-related changes in fundamental processes controlling gene expression in human muscle.

## Results

### Methodology overview

We describe our study design in **Fig 1**.

**Figure 1.**
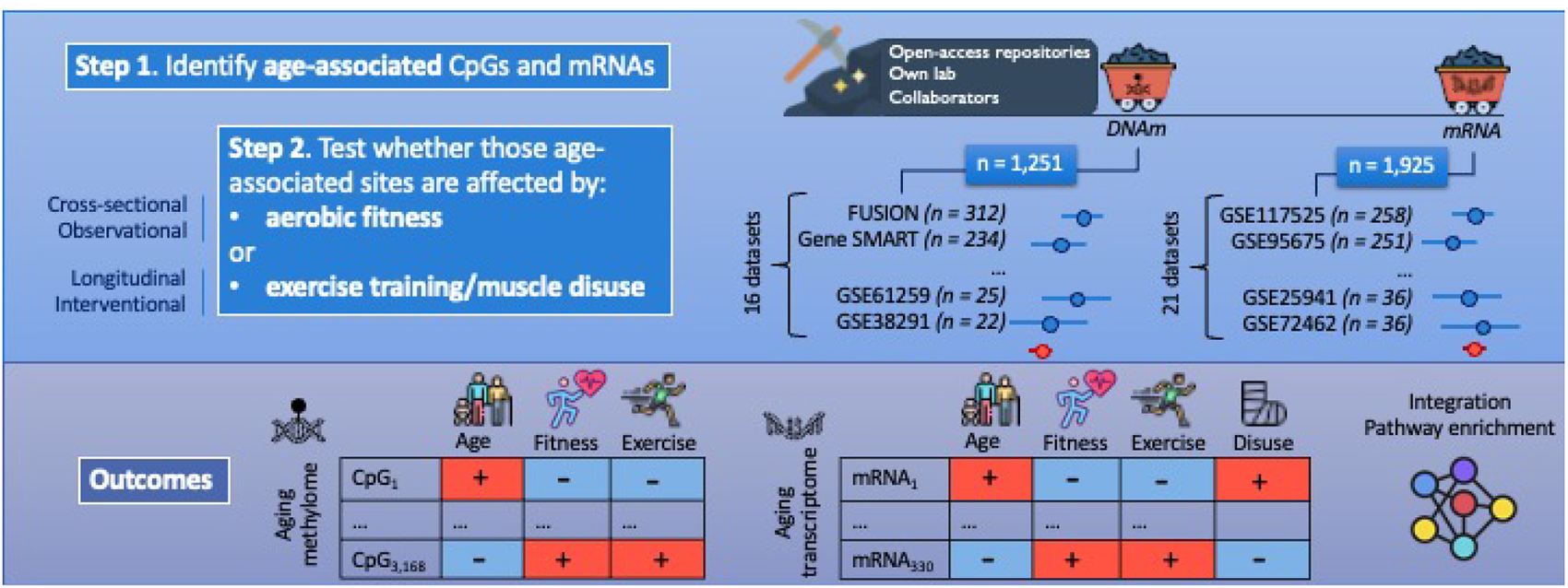
Study overview. First, we performed a large-scale data mining exercise to gather all existing DNA methylation (DNAm) and mRNA expression microarray datasets from our own lab, our network of collaborators, and open-access repositories (GEO, dbGAP, ArrayExpress). See Methods for inclusion criteria of datasets. Step 1: we identified age-related changes in DNAm and mRNA expression in human skeletal muscle by meta-analysing 1,251 samples across 16 cohorts (DNAm) and 1,925 samples across 21 cohorts (mRNA). Step 2: first, we performed a cross-sectional association between DNAm levels at age-related CpGs or expression levels at age-related mRNAs, and VO2max. Then, we determined whether DNAm levels at age-related CpGs, and expression levels at age-related mRNAs changed after exercise training, and we assessed whether expression levels at age-related mRNAs changed after muscle disuse. Finally, we performed a series of OMIC integrations and pathway analyses to identify the molecular pathways affected by age, VO2max, exercise training, and/or muscle disuse across both OMIC layers. <u>Note</u>: we only had OMIC data available at the transcriptomic level following muscle disuse. <u>Note 2</u>: summary statistics for the exercise- and disuse-induced mRNA changes came from two recent meta-analyses^24,25^.

First, we identified the DNAm loci and transcripts that change with age in human skeletal muscle, by systematically mining and analysing DNAm and mRNA microarray data from our laboratory, online databases, and our collaborators’ laboratories. We conducted a random-effects epigenome-wide association study (EWAS) meta-analysis of age across 1,251 samples from 16 independent cohorts (Supplementary Table 1), and a random-effects transcriptome-wide association study (TWAS) meta-analysis of age across 1,925 human muscle samples from 21 independent cohorts (Supplementary Table 2). To uncover the effect of aerobic fitness and exercise training on the aging methylome and transcriptome of skeletal muscle, we then restricted our analysis to the identified age-related Differentially Methylated Positions (DMPs) and Differentially Expressed Genes (DEGs).

We first assessed whether an objective gold-standard measure of aerobic fitness (VO_2max_) was associated with younger methylome and transcriptome profiles in skeletal muscle (**Fig 1**). At the epigenetic level, we conducted a random-effects meta-analysis of VO_2max_ across 439 samples from 5 independent cohorts; at the transcriptomic level, we conducted a random-effects meta-analysis of VO_2max_ across 354 samples from 5 independent cohorts (**Table 1**). There was a reasonably large range of VO_2max_ in each of these cohorts (SD > 5 mL/min/kg, **Table 1**), which is essential to detect differences in OMIC aging between individuals with variable fitness levels.

**Table 1.** Characteristics of the cohorts used in the cross-sectional analysis (association between age-related CpGs or mRNAs and maximal oxygen uptake (VO2max)).

| | Cohort | Number of samples | Number of unique individuals | Age (years) | Sex (% male) | $VO_{2max}$ (mL/min/kg) | Data availability |
| --- | --- | --- | --- | --- | --- | --- | --- |
| DNA methylation | Gene SMART<sup>9,10</sup> | 234 | 66 | $32 \pm 8.1$ | 80% | $48 \pm 9.0$ | GSE151407, GSE171140 |
| | Finnish Twin Cohort<sup>17</sup> | 73 | 73 | $43 \pm 16$ | 38% | $36 \pm 8.5$ | Per request |
| | E-MTAB-11282<sup>25</sup> | 52 | 13 | $45 \pm 11$ | 38% | $23 \pm 5.0$ | E-MTAB-11282 |
| | EXACT | 48 | 16 | $33 \pm 10$ | 100% | $42 \pm 7.8$ | Per request |
| | CAUSE<sup>22</sup> | 32 | 16 | $60 \pm 5.3$ | 0% | $30 \pm 6.1$ | GSE213029 |
| mRNA expression | GSE18732<sup>26</sup> | 117 | 117 | $54 \pm 11$ | 69% | $28 \pm 9.6$ | GSE18732 |
| | HERITAGE<sup>27,28</sup> | 82 | 42 | $34 \pm 14$ | 58% | $36 \pm 9.0$ | GSE117070 |
| | GSE44818<sup>29</sup> | 72 | 12 | $30 \pm 7.2$ | 100% | $60 \pm 6.0$ | GSE44818 |
| | The Malmö Prevention Study<sup>30</sup> | 43 | 43 | $66 \pm 1.5$ | 100% | $28 \pm 6.4$ | E-CBIL-30 |
| | MSAT<sup>31</sup> | 39 | 39 | $36 \pm 8.3$ | 100% | $52 \pm 8.1$ | Per request |

As cross-sectional analyses can be confounded by unmeasured factors (e.g. lifelong dietary patterns, socioeconomic status), we tested directly whether an exercise training program could shift OMIC profiles towards a younger state, in an interventional, longitudinal setting. We conducted a random-effects meta-analysis of DNAm changes following aerobic, high-intensity interval, or resistance training across 401 samples from 6 independent exercise training interventions (**Fig 1** **Table 2**); we also extracted summary statistics at age-related DEGs from a published meta-analysis of transcriptomic changes induced by exercise training^32^. Finally, and to further support the causal effect of exercise training in the shift of muscle OMIC patterns towards younger profiles, we tested whether age-related transcriptomic profiles were altered following a *decrease* in physical activity; we extracted summary statistics at age-related DEGs from a published meta-analysis following forced immobilisation protocols33.

**Table 2.** Participant characteristics from the cohorts in the interventional analysis (changes in levels of age-related CpGs or mRNA after exercise training or muscle disuse).

|  | Cohort | Number of samples | Number of unique individuals | Age (years) | Sex (% male) | Intervention | Dataset access |
| --- | --- | --- | --- | --- | --- | --- | --- |
| DNA methylation | Gene SMART<sup>9,10</sup> | 196 | 66 | $32 \pm 8.1$ | 80% | 4 weeks of HIIT repeated twice after a > 1 year washout, and an additional 8 weeks of HIIT | GSE151407, GSE171140 |
| | E-MTAB-11282<sup>25</sup> | 52 | 13 | $45 \pm 11$ | 38% | 8 weeks of aerobic training | E-MTAB-11282 |
|  | EPIK<sup>21</sup> | 48 | 14 | 45 ± 22 | 100% | 12 weeks of resistance training repeated twice after a 12-week washout (older individuals) or 2-week immobilization (younger individuals) | Per request |
|  | GSE114763<sup>34</sup> | 39 | 8 | 29 ± 6 | 100% | 7 weeks of resistance training repeated twice after a 7-week washout | GSE114763 |
|  | EpiTrain<sup>35</sup> | 34 | 17 | 27 ± 4 | 44% | 3 months of aerobic training | GSE60655 |
|  | CAUSE<sup>22</sup> | 32 | 16 | 60 ± 5.3 | 0% | 5 months of aerobic training | GSE213029 |
| mRNA expression | ExTraMeta<sup>32</sup> | 952 | 476 | Meta-analysis of 28 datasets |  | Exercise training of various types and duration | https://www.extrameta.org |
|  | MetaMEx<sup>33</sup> | 250 | 125 | Meta-analysis of 7 datasets |  | Forced immobilisation of various durations | https://metamex.serve.scilifelab.se/ |

#### Skeletal muscle aging alters DNA methylation and expression of genes involved in muscle structure and metabolism

Age-related changes to the methylome are widespread yet small (typically ∼1% change in methylation per decade of age); out of the 595,541 tested CpG sites, we identified 3,168 differentially methylated positions (DMPs) associated with age at FDR < 0.005^36^ (Supplementary Table 3), 73% of which were hypomethylated (Supplementary Fig 1A). This is in concordance with our previous findings^10^, and in line with recent findings at the single-cell level^37^. While DMPs were not enriched in any particular canonical pathway or expression signatures of genetic and chemical perturbations, they were overrepresented in two gene ontology terms related to muscle structure (contractile fibre, I band; Supplementary Fig 1B), and in two human phenotype ontologies entirely consistent with musculoskeletal aging (“difficulty climbing stairs”, and “muscle weakness”; Supplementary Fig 1B).

**Supplementary Figure 1.**
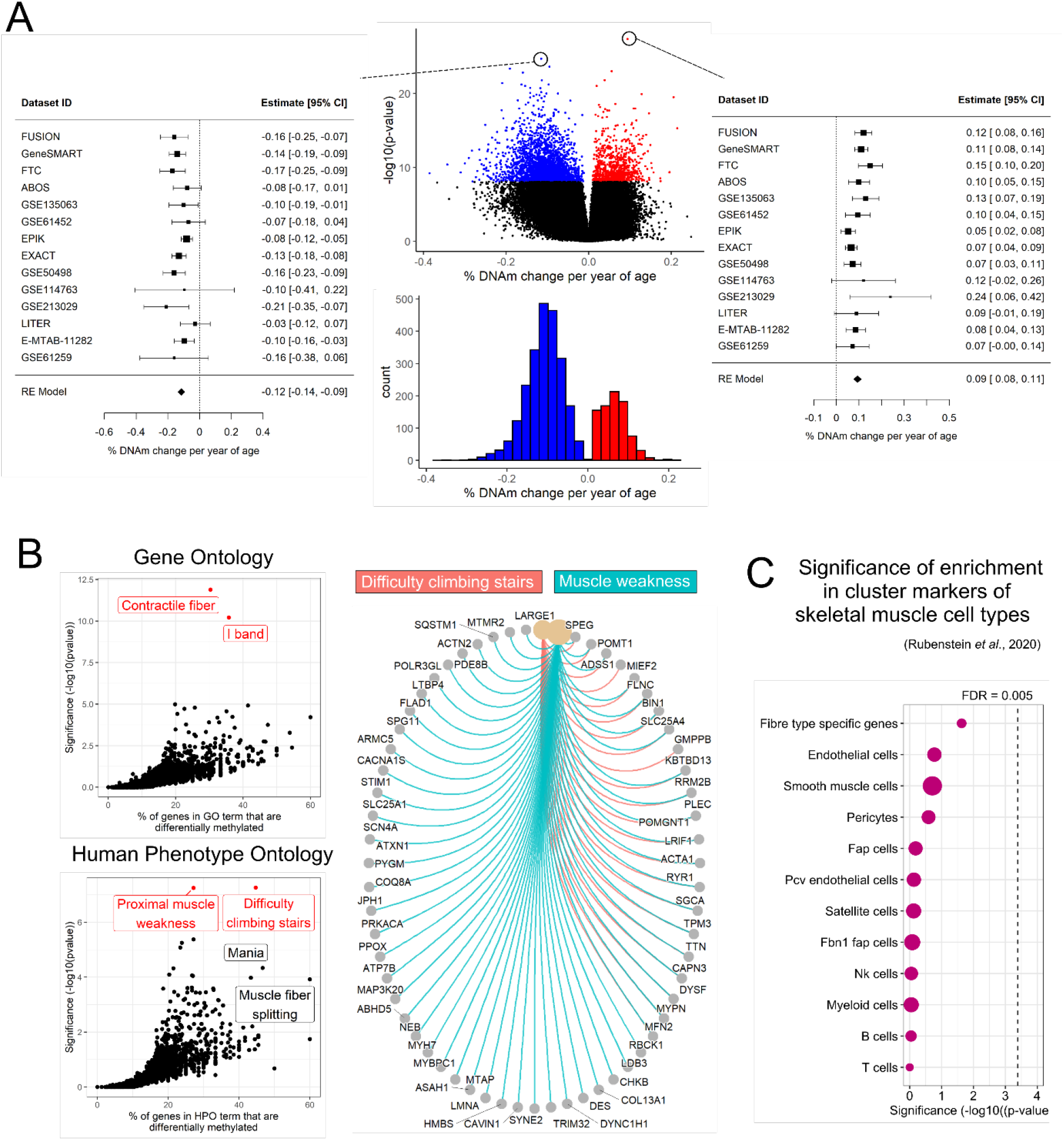
EWAS meta-analysis of age reveals small, widespread DNA methylation changes at genes encoding structural components of muscle. A) Volcano plot of the effect size (% DNAm change per year of age) and significance of age for each CpG included in the meta-analysis (pooled results of 16 independent muscle datasets); colored dots are CpGs that are differentially methylated at a false discovery rate (FDR) < 0.005; we highlighted the top hypo-(left) and hyper-(right) methylated CpG in forest plots to show the estimates and confidence intervals in each dataset. B) Overrepresentation analysis of the age-related differentially methylated positions for Gene Ontology, and Human Phenotype Ontology (HPO) from MSigDB; pathways in red are significant at an FDR < 0.005. The circular plot displays genes from the two significant HPOs (“difficulty climbing stairs”, and “muscle weakness”) that showed age-related alterations at the DNAm level. C) Dotplot of the enrichment of DMPs in marker genes for 12 cell types identified in a single-cell sequencing study of human skeletal muscle38.

Out of the 16,657 tested transcripts, we identified 330 differentially expressed genes (DEGs) at FDR < 0.005 (Supplementary Table 4), 68% of which were downregulated with older age (Supplementary Fig 2A). While DEGs were not enriched in any particular canonical pathway, they were overrepresented in several gene ontology terms related to mitochondrial function and energy production (ADP and ATP metabolic processes, generation of precursor metabolites and energy, energy derivation by oxidation of organic compounds, mitochondrial protein containing complex, organelle inner membrane; Supplementary Fig 2B). Furthermore, they were also overrepresented in expression signatures of genetic and chemical perturbations (incl. genes up-regulated in differentiating myoblasts upon expression of *PPARGC1A*^39^, genes differentially regulated in myoblasts with *IGF2BP2* knockdown^40^, and genes that comprise the mitochondria gene module41). DEGs were also enriched in two human phenotype ontologies related to muscle function (“exercise intolerance”, and “myoglobinuria”; Supplementary Fig 1B).

Finally, we estimated whether the age-related DNAm and mRNA changes were possibly driven by changes in cell type composition. We tested whether the DMPs and DEGs were overrepresented in twelve gene sets containing curated cluster markers for cell types identified in a single-cell sequencing study of human skeletal muscle^38^. We found no significant enrichment of DMPs in any of the cell type marker genes (Supplementary Fig 1C), suggesting that the age-related DNAm changes are not confounded by changes in cell type proportions. In contrast, DEGs were enriched for genes whose expression differs between type I and type IIa fibres (Supplementary Fig 2C), with a change in mRNA expression suggestive of an increase in type I fibre % with older age; *TPM3, PDLIM1, MYOZ2* are all markers of type I fibres and increased in expression with age, while *PKM, ENO3 and PFKM* are all markers of type IIa fibres and decreased in expression with age. DEGs also showed a trend for enrichment in marker genes of smooth muscle cells that make up the walls of blood vessels (Supplementary Fig 2C), but it is unclear whether it reflected an increase or a decrease in the proportion of smooth muscle cells, as the signal was inconsistent: 17/28 of the marker genes increased in expression, and 11/28 decreased in expression.

**Supplementary Figure 2.**
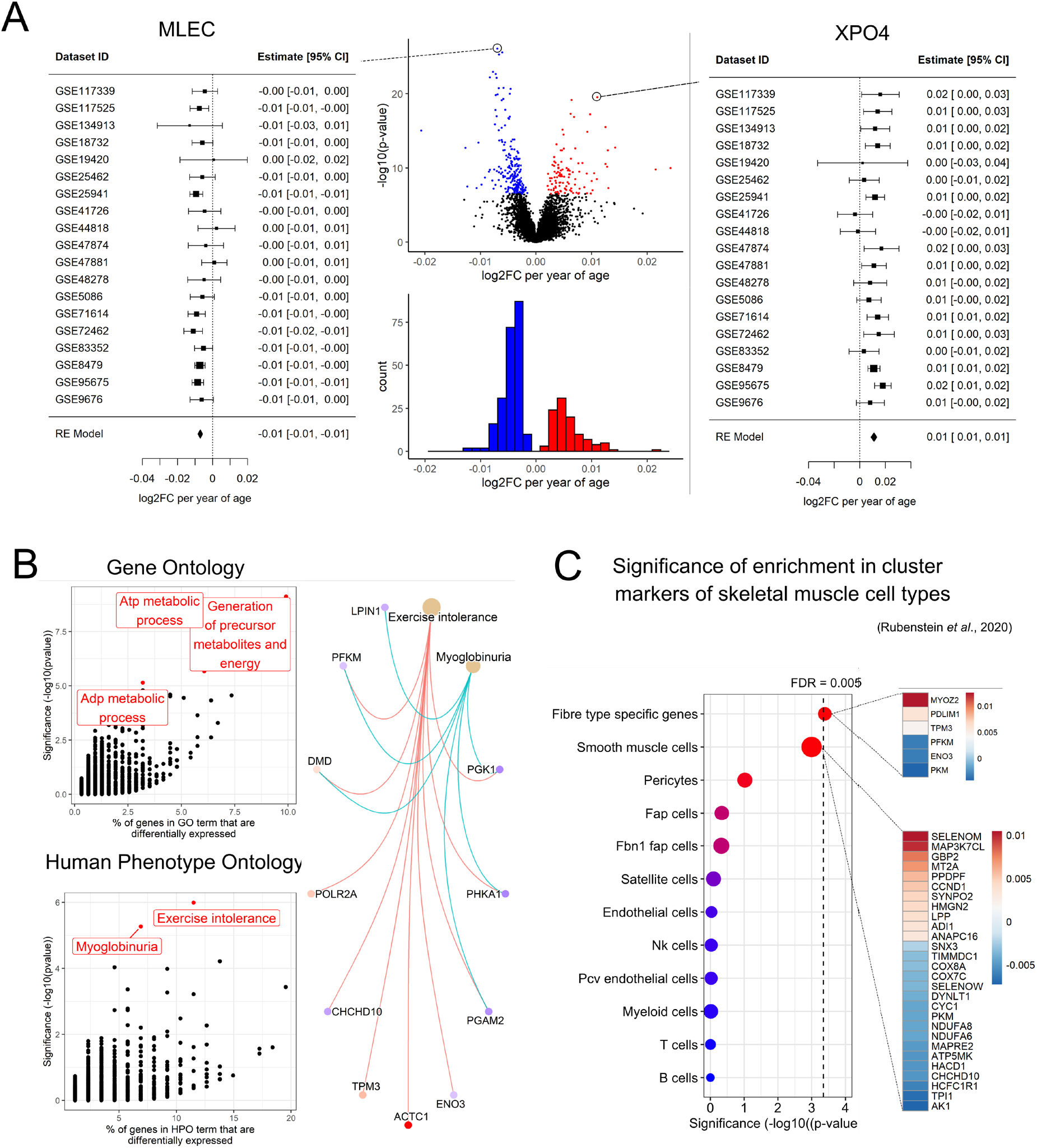
TWAS meta-analysis of age reveals mRNA expression changes at genes involved in mitochondrial function and energy production. A) Volcano plot of the magnitude (log2 fold-change (log2FC) per year of age) and significance of age effects for each gene included in the meta-analysis (pooled results of 21 independent muscle transcriptomic datasets); coloured dots are genes that are differentially expressed at a false discovery rate (FDR) < 0.005; we highlighted the top down-(left) and up-(right) regulated genes in forest plots to show the estimates and confidence intervals in each dataset. B) Overrepresentation analysis of the age-related differentially expressed genes (DEGs) for Gene Ontology (GO), and Human Phenotype Ontology (HPO) from MSigDB; pathways in red are significant at an FDR < 0.005. The circular plot displays DEGs from the two significant HPOs (“exercise intolerance”, and “myoglobinuria”). C) Dotplot of the enrichment of DEGs in marker genes for 11 cell types identified in a single-cell sequencing study of human skeletal muscle38. The heatmap displays the log2FC of the DEGs that are marker genes for smooth muscle cells.

As the DMPs and DEGs were associated with related yet distinct ontologies, we further examined the overlap between differentially methylated genes (DMGs) and DEGs. We identified 63 genes that were altered both at the epigenetic and transcriptional levels during aging (Supplementary Fig 3A, Supplementary Table 5). This overlap between age-related changes at the epigenetic and transcriptomic levels was greater than expected by chance alone, as DMGs were more likely to also be DEGs than non-DMGs (p-value for over-representation = 0.00011, Supplementary Fig 3B); conversely, DEGs were more likely to be DMGs than non-DEGs (p-value for over-representation = 1.7 × 10^−5^, Supplementary Fig 3B).

**Supplementary Figure 3.**
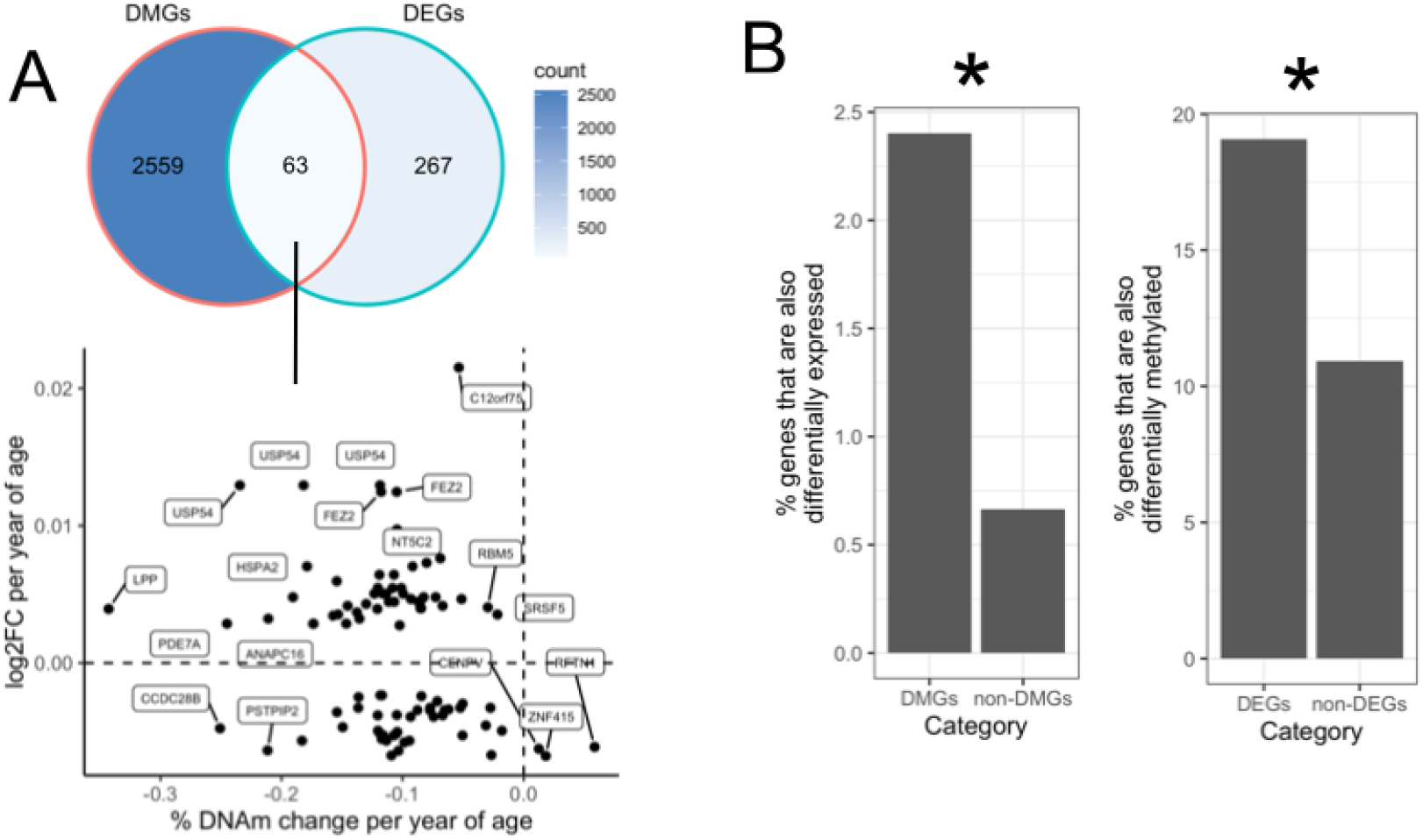
Overlap between age-related differentially methylated genes (DMGs) and differentially expressed genes (DEGs). A) The 3,168 DMPs were located in 2,622 unique genes, 63 of which were also DEGs. On the bottom graph, each dot represents a DMP-DEG pair, with the change in DNAm level (x-axis) and mRNA level (y-axis) per year of age. Note that some genes appear multiple times on the plot as multiple DMPs could be annotated to the same gene (e.g. FEZ2, USP54). B) Left, proportion of genes within DEGs and non-DEGs that are also differentially methylated with age; right, proportion of genes within DMGs and non-DMGs that are also differentially expressed with age. *p < 0.005 in over-representation test (Fisher’s exact test).

##### CRF and exercise training are associated with younger epigenetic and transcriptomic profiles in human skeletal muscle, in contrast to muscle disuse

We identified 25 age-related DMPs and one age-related DEGs that showed a significant association with VO_2max_ (FDR < 0.005, Supplementary Tables 3 & 4), but it is likely that we were underpowered to detect more associations at a high level of confidence. A quantile-quantile plot of p-values for the association between VO_2max_ and DNAm or mRNA levels at age-related DMPs or DEGs, showed a clear increase (inflation) above the diagonal line (Supplementary Fig 4A & B). This diagonal line indicates the expected p-value distribution under the assumption (null hypothesis) that the p-values follow a uniform [0,1] distribution (i.e. that there are no true associations between VO_2max_ and DNAm or mRNA levels at age-related DMPs and DEGs). The amount of departure from this diagonal line correlates with the expected number of true associations^42^. We did not identify age-related DMPs that were significantly altered following exercise training at FDR < 0.005, but with substantially more samples (n = 952 from 28 datasets) and therefore statistical power at the transcriptional level, we found 40 age-related DEGs that were altered following exercise training (Supplementary Table 3). Importantly, at age-related DMPs and DEGs most significantly associated with VO_2max_ or altered following exercise training (the points on the far right of the Q-Q plots), VO_2max_ and training were associated with changes that directly countered the effect of age (Supplementary Fig 4, C & D). In other words, DMPs or DEGs whose methylation or expression levels decreased with age overwhelmingly increased in methylation or expression with higher VO_2max_ and following exercise training; DMPs or DEGs whose methylation or expression levels increased with age overwhelmingly decreased in methylation or expression with higher VO_2max_ and following exercise training. Finally, we observed a very pronounced effect of muscle disuse on the aging transcriptome, despite a substantially lower sample size (n = 250 from 7 datasets, **Table 2**): 68 age-related DEGs were significantly altered following forced immobilisation at FDR < 0.005 (Supplementary Table 4, Supplementary Fig 4E). DEGs that were downregulated with age tended to decrease in expression following muscle disuse, and vice versa.

**Supplementary Figure 4.**
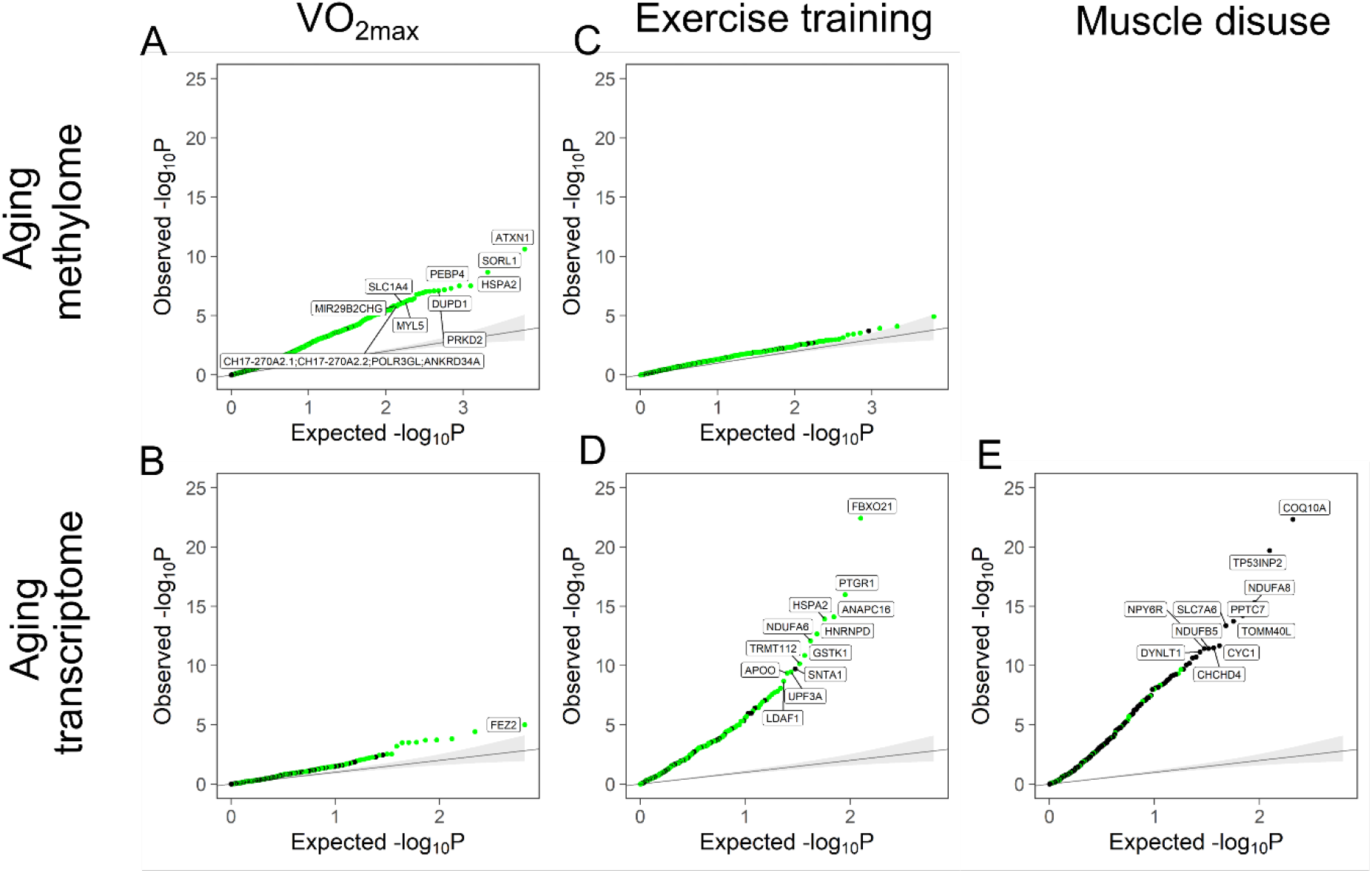
Quantile-quantile plot of p-values for the effect of VO2max, exercise training and muscle disuse on the aging methylome and transcriptome. The observed p-values for each age-related DMP or DMG are sorted from largest to smallest and plotted against expected values from a theoretical χ2-distribution. If some observed p-values are clearly more significant than expected under the null hypothesis, points will move towards the y-axis; green points represent DMPs or DEGs for which the effect of age contrasts with the effect of VO2max or exercise or disuse, while black points represent DMPs or DEGs for which the effect of age is in line with the effect of VO2max or exercise or disuse.

The contrast between age-related and VO_2max_-and exercise-related changes are visible when all age-related DMPs and DEGs were taken into account; we noted strong negative correlations between the effects of age and VO_2max_ across all age-related DMPs (**Fig 2A**, Spearman correlation ρ = -0.39, *p* < 2.2 × 10^−16^, **Fig 2B**), and across all age-related DEGs (**Fig 3A**, Spearman correlation ρ = -0.18, *p* = 0.0013, **Fig 3B**), as well as strong negative correlations between the effects of age and exercise training over all age-related DMPs (**Fig 2A**, Spearman correlation ρ = -0.37, *p* < 2.2 × 10^−16^) and across all age-related DEGs (**Fig 3A**, Spearman correlation ρ = – 0.38, *p* = 6.7 × 10^−12^). The “pro-aging” effect of muscle disuse was visible at the scale of the whole aging transcriptome; we found a strong positive correlation between the effects of age and immobilisation over all age-related DEGs (**Fig 3A**, Spearman correlation ρ = 0.43, *p* < 2.2 × 10^−16^).

**Figure 2.**
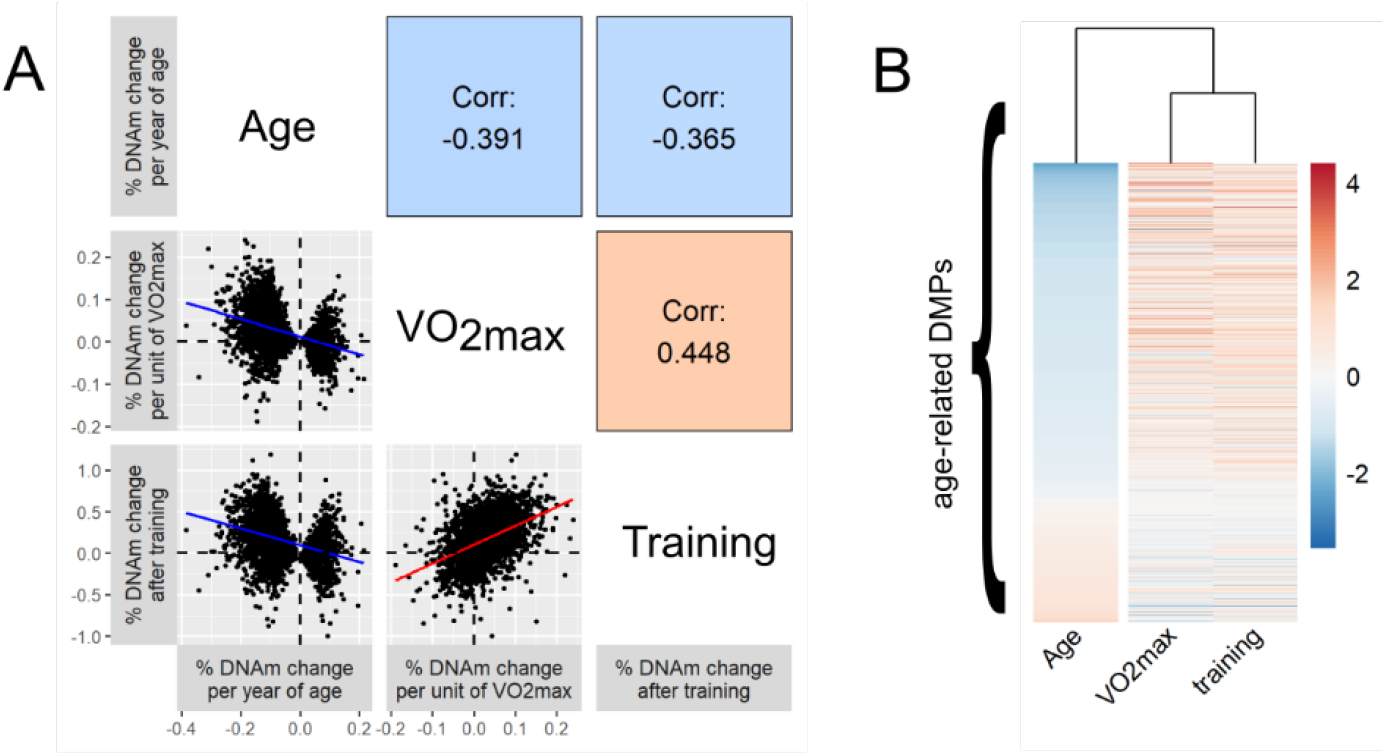
Aerobic fitness and exercise training have similar associations with muscle epigenetic profiles, which contrast with those seen with age. A) Pairwise correlation (Spearman) between the effect sizes of age, aerobic fitness (VO2max) and exercise training at age-related DMPs. Each dot corresponds to one of the 3,168 age-related DMPs, and the axes represent the magnitude of effect for age, VO2max and exercise training. B) Unsupervised hierarchical clustering of the effect sizes of age, VO2max and exercise training at age-related DMPs (ordered from the most hypomethylated to most hypermethylated with age). Note that the legend is arbitrary as effect sizes were scaled to an SD of 1 for age, VO2max and exercise training to be comparable.

**Figure 3.**
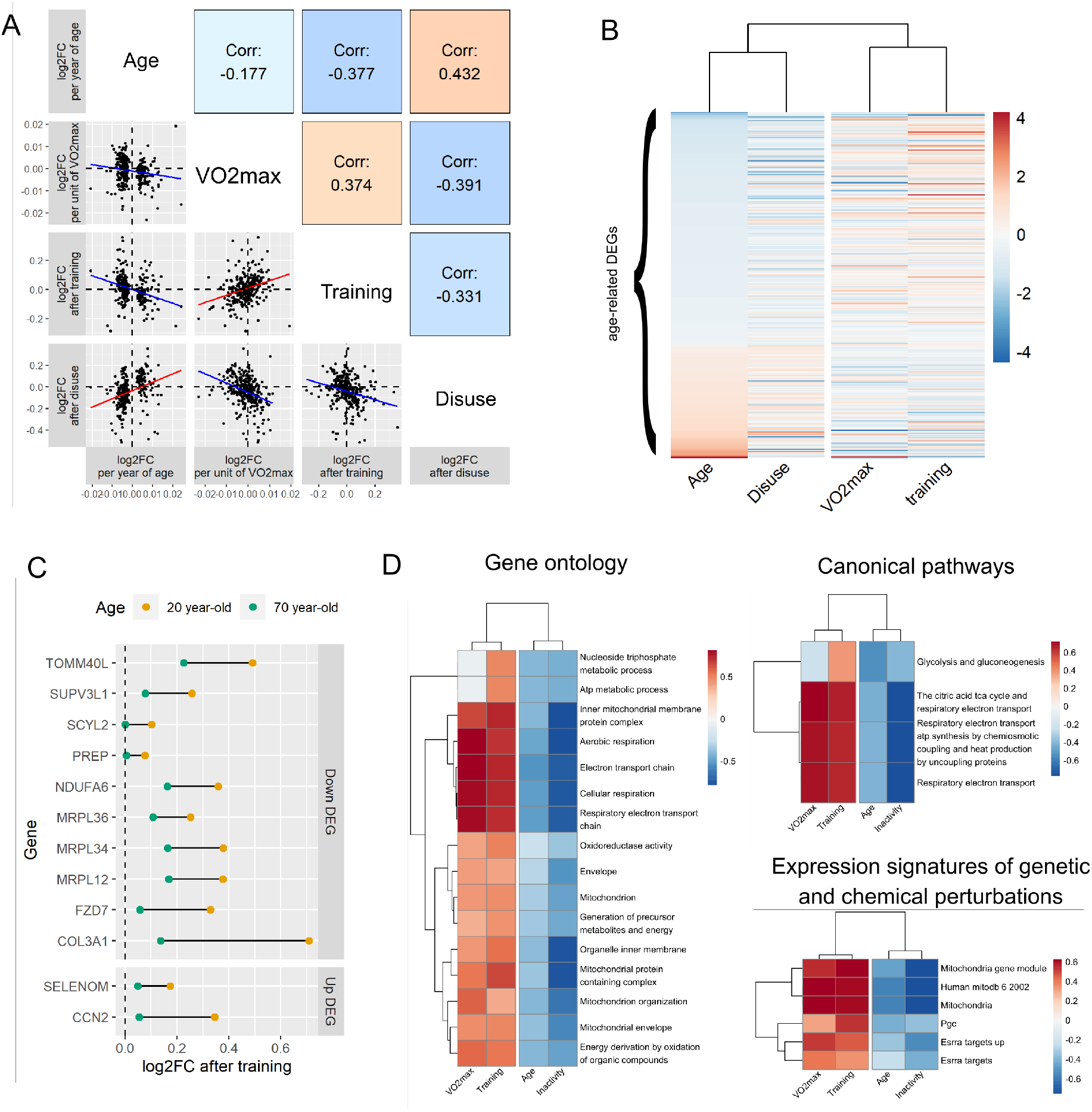
Aerobic fitness and exercise training show similar associations with muscle transcriptomic profiles, and contrast with those seen with age and muscle disuse. A) Pairwise correlation (Spearman) between the effect sizes of age, aerobic fitness (VO2max), exercise training and muscle disuse at age-related DEGs. Each dot corresponds to one of the 330 age-related DEGs, and the axes represent the magnitude of effect for age, VO2max, exercise training and disuse. B) Unsupervised hierarchical clustering of the effect sizes of age, VO2max, exercise training and disuse at age-related DEGs (ordered from the most downregulated to most upregulated with age). Note that the legend is arbitrary as effect sizes were scaled to an SD of 1 for age, VO2max, exercise training and disuse to be comparable. C) Comparison of the magnitude of exercise-induced changes in gene expression between a hypothetical 20 year-old and 70 year-old. The genes displayed are those for which “age” was a significant moderator according to the meta-regression conducted by Amar et al.32 “Down DEG” = gene whose expression decreases during normal aging; “Up DEG” = gene whose expression increases during normal aging. D) Multi-contrast enrichment comparing the effects of age, VO2max, exercise training and muscle disuse at age-related DEGs. Genes related to mitochondrial function showed clear indications of downregulation during aging and following muscle disuse, while being simultaneously upregulated with higher VO2max and following exercise training. This was visible across the Gene Ontology, Canonical Pathways, and Expression Signature of Genetic & Chemical Perturbations gene sets.

To further highlight the association between VO_2max_ and muscle OMIC aging, we performed principal component analysis (PCA) for DMPs in the Gene SMART cohort, and for DEGs in the GSE18732 cohort. These two cohorts have the largest sample size and VO_2max_ range in our study (**Table 2**). In the Gene SMART cohort, individuals clustered by VO_2max_ levels both on Dimension 1 and Dimension 3 (Pearson correlation *p* = 0.0011 for PC1 and *p* = 0.0047 for PC3), indicating that individuals of similar fitness levels show similar patterns of DNAm at age-related DMPs (**Fig 4A**). Similarly, in the GSE18732 cohort, individuals tended to cluster by VO_2max_ levels both on Dimension 1 and Dimension 3 (Pearson correlation *p* = 0.05 for PC1 and *p* = 0.0019 for PC3), indicating that individuals of similar fitness levels have similar mRNA levels at age-related DEGs (**Fig 4B**).

**Figure 4.**
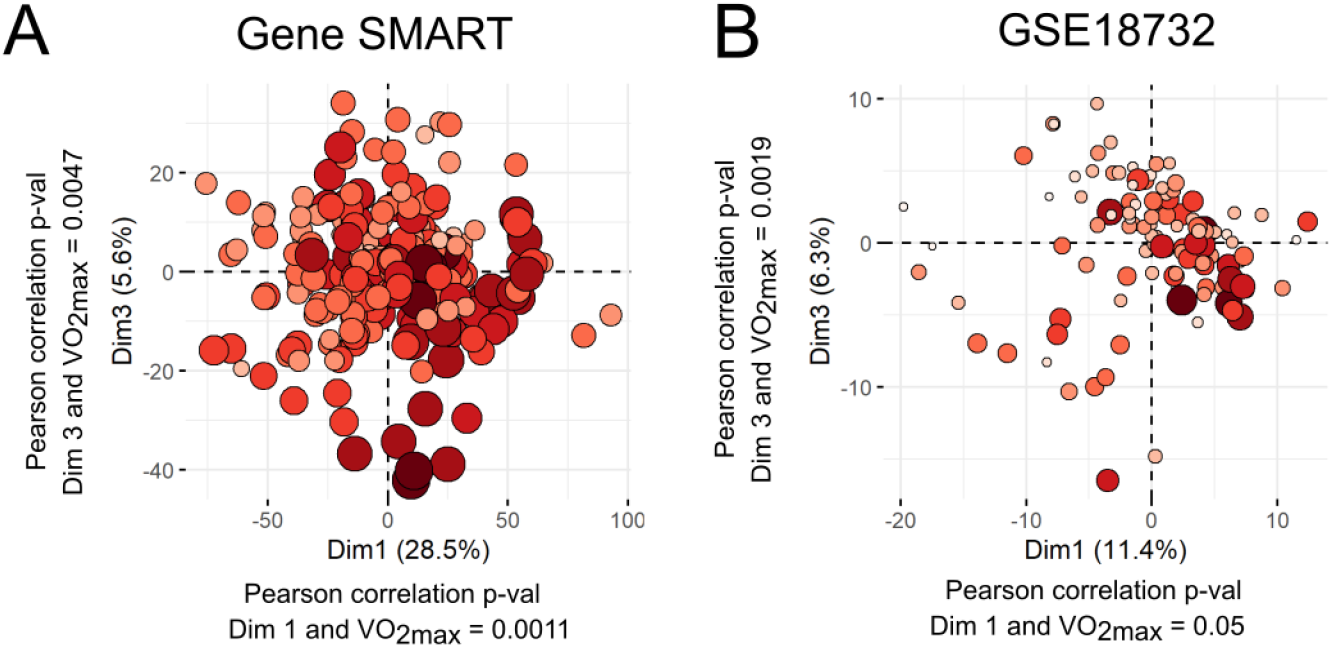
Principal component analysis of individuals from the Gene SMART cohort at age-related DMPs, and individuals from the GSE18732 cohort at age-related DEGs. We used principal component analysis (PCA) to reduce dimensionality and show each individual on a two-dimensional graph. Individuals from the Gene SMART cohort (A) and from the GSE18732 cohort (B) are colored according to their baseline VO2max levels. Lower levels of VO2max are indicated by smaller circles in lighter colors, while higher levels of are indicated by larger circles in darker reds. To objectively test the clustering of individuals according to VO2max, we ran Pearson correlations between individual coordinates on Dimension 1, Dimension 2 or Dimension 3 and VO2max.

##### Older individuals have a blunted response to exercise at a small fraction of age-related DEGs

We then explored whether the effects described above were applicable to the entire lifespan (i.e. whether older individuals were as able as young individuals to reap the “anti-aging” benefits of exercise training). We could not test this hypothesis at the DNAm level due to the more limited number of older individuals with DNAm data in the studied cohorts (**Tables 1 & 2**), but we investigated this in the transcriptional response to exercise training as effect sizes came from 28 different cohorts with a wide variability in age range (Table 2). We extracted summary statistics from the original paper from Amar *et al*.^32^, and identified 12 DEGs (3% of all DEGs) whose exercise-induced change in expression levels depends on age. For all 12 DEGs, older individuals showed a blunted response to exercise training (**Fig 3C**).

##### Mitochondrial and metabolic pathways are simultaneously inhibited by age and muscle disuse, and enhanced with greater CRF and by exercise training

We compared the associations between age, aerobic fitness, exercise training, and muscle disuse in an integrated, multi-contrast enrichment analysis that uses a rank-MANOVA based statistical approach43. We could only perform this multi-contrast analysis at the transcriptional level, as this statistical approach has not been adapted for DNAm data (i.e. it has not been optimised to take into account the severe bias in gene-set analysis applied to genome-wide methylation data^44,45^). This multi-contrast analysis identified mitochondrial and metabolic pathways as simultaneously inhibited by age and muscle disuse, while enhanced by aerobic fitness and exercise training (**Fig 3D**). The effects of VO_2max_ and exercise training were highly consistent both at the epigenetic and transcriptomic levels (**Fig 2** **& 3**). Conversely, the effect of muscle disuse contrasted with both VO_2max_ and exercise training (**Fig 2** **& 3**).

## Discussion

We showed, in a large sample (> 3,200) of human muscles, that higher aerobic fitness is associated with younger epigenetic and transcriptomic profiles. In line with this, exercise training shifts the aging muscle epigenome and transcriptome towards a younger profile, while muscle disuse significantly accelerates transcriptomic aging. The magnitude of “anti-aging” or “aging” effects were highly site- and gene-dependent, suggesting that exercise training can reverse specific OMIC changes occurring during normal aging in human skeletal muscle. Transcriptomic integration revealed a pronounced deterioration of mitochondrial function and energy production during aging, which is accelerated by forced immobilisation but restored by exercise training.

Although there were few age-related DMPs and DMGs significantly associated with CRF or exercise training, a close inspection of all statistical tests performed suggested that this was more likely due to a lack of statistical power than an absence of true associations. In support of this, we observed a striking similarity between the effect sizes of aerobic fitness and exercise training on the aging muscle methylome and transcriptome, which directly contrasted with the effects of age and forced immobilisation. Adults lose about ∼2.5-3 mL/min/kg of VO_2max_ per decade of age^46^, partly because of primary aging (i.e. the inevitable deterioration of cellular structure and biological function, independent of disease or harmful lifestyle or environmental factors^47^), but also due to a decline in physical activity levels as we age^48^. Therefore, the anti-aging effect of exercise and the aging effect of disuse are perhaps unsurprising, given that some of the age-related signal captured in the EWAS and TWAS meta-analysis of age reflect a decline in physical activity levels, rather than primary aging per se. This exemplifies the complex nature of aging biology, and highlights the challenge of assessing which factors mutually affect each other when causality becomes circular^49^ (e.g. aging leads to a decline in physical activity/fitness levels, and a decline in physical activity/fitness levels leads to aging^50^). In a small study of older men, master athletes showed hypomethylation in the promoter of genes involved in energy metabolism and muscle structure, compared with men who reported lifelong sedentary behaviour51. However, without a young control group, this study could not distinguish the age-related changes that can be counteracted by exercise, from those that remain unaffected by even the most extreme exercise regimes. Without adjusting for other environmental confounders, it is also unclear whether these effects come from the exercise regime or rather from the effects of other lifestyle factors that correlate strongly with high physical activity levels (e.g. a healthy diet). In our study, the interventional data (i.e. exercise training & muscle disuse protocols) entirely supported the cross-sectional findings (i.e. associations with CRF), suggesting that the association between CRF and OMIC aging is due to exercise training rather than other unmeasured confounding factor.

We used a large-scale data mining approach to achieve an unprecedented sample size (> 1,200 epigenetic profiles and >1,900 transcriptomic profiles across 37 cohorts). We applied random-effects meta-analyses allowing the effects of age, fitness, exercise and disuse to vary between cohorts while maintaining the specificity of each dataset (i.e. we did not force a normalisation across datasets). Our results may therefore be applicable to a broad range of individuals (sex, health/training status) and exercise regimes (training type/duration). However, the cohorts profiled for DNAm included a majority of healthy, young/middle-aged, male individuals, so we cannot confidently extrapolate the DNAm results to all populations. It is possible that older, or diseased individuals show blunted responses to exercise training, or that specific training regimes (e.g. endurance vs resistance training) lead to anti-aging effects that are entirely specific to that training regime. We could only test this at the transcriptional level, and we found that a few age-related DEGs showed a blunted response to exercise training in older individuals.

While we observed the anti-aging effect of exercise training at both the epigenetic and transcriptomic levels, and despite a significant overlap between DMPs and DEGs, the molecular pathways affected by age were distinct between the two OMIC layers. It may be due to differences in gene coverage between the DNAm and mRNA arrays, or it could reflect differences in aging mechanisms at the epigenetic and transcriptomic levels. Nevertheless, the integration of all effects at the transcriptomic level clearly showed a downregulation of mitochondrial and energy metabolism pathways during aging and following muscle disuse, which was restored by aerobic fitness and exercise training. This is in line with the known beneficial effect of exercise on mitochondrial function52. The integration was not feasible at the DNAm level because a given CpG can be annotated to multiple genes, and a given gene can harbour multiple CpGs, severely biasing the statistical test for enrichment^44,45^.

A major challenge in OMIC analysis is determining whether changes are due to a modification of the intrinsic profiles of the cells, or to a change in the relative proportions of different cell types in the sample. We could not directly estimate the proportions of different cell types in our samples, as deconvolution algorithms have currently not been developed for human skeletal muscle, whether at the methylation^53^ or transcriptional level. We therefore tested for an enrichment of DMGs and DEGs in genes whose expression differ between muscle cell types^38^. While we found no evidence of confounding by cell type in the epigenetic analysis, some of the age-related changes in mRNA were indicative of an increase in the proportion of type I fibres (vs type II fibres). This is surprising given that age does not affect the relative proportion of different fibre types, but rather the size and distribution of fibres within the muscle^54^. Furthermore, we observed younger OMIC patterns in individuals of higher aerobic fitness levels, yet fitter individuals typically harbour greater proportions of type I fibres^55^. Therefore, it is likely that the DNAm and mRNA expression changes were intrinsic to muscle cells, rather than reflecting a shift in the proportions of different cell types within the muscle. To answer this question, future studies should investigate the effects of age, aerobic fitness, exercise training and muscle disuse on OMIC profiles within individual cell types using cell sorting, or single-cell methods.

We avoided using epigenetic clocks in this analysis for multiple reasons. There are only two clocks currently available that could be applied to muscle DNAm data, namely the Horvath pan-tissue clock^56^ and the MEAT clock we recently developed for human muscle9 (all other clocks were developed for non-muscle tissue). First, both the pan-tissue and MEAT clocks were trained to predict *chronological* age, which is a poor proxy for clinically relevant measures of biological age (this has been highlighted by others^8,57,58^ and is the reason for the development of second-generation clocks, such as PhenoAge^59^ and GrimAge^60,61^, clocks better adapted to longitudinal data such as DunedinPoAm^62^ and DunedinPACE^63,^ and clocks able to disentangle damaging and adaptive changes during aging^64^). There are currently too few DNAm datasets with corresponding measurements of muscle function (e.g. mitochondrial function, contractile properties, etc.) to develop an epigenetic clock that would capture muscle biological age. Second, the pan-tissue clock is poorly calibrated in skeletal muscle, and most of the muscle datasets from the present study were used to generate the MEAT clock^9^. This means that epigenetic age estimations using the MEAT clock would be severely biased and unsuitable to assess the effects of fitness and exercise on epigenetic aging. Finally, until recently^65^, epigenetic clocks only selected a limited number of CpGs to maximise prediction accuracy, which means they would discard information at many potentially relevant CpGs associated with age. We adopted a broader perspective to look at the entire aging methylome and examine the effect of exercise training on this aging trend. We were unable to assess the functional effects of the age-related OMIC changes on muscle structure, function, and metabolism, so we cannot firmly conclude that the effect of exercise training led to gains in muscle function or quality. Future studies combining OMIC profiles with genetic data to implement Mendelian Randomization analyses64 that would determine whether OMIC aging leads to a decline in muscle function, and whether the beneficial effects of fitness and exercise training on muscle function are mediated by a reversal of OMIC aging.

In conclusion, using an unprecedented number of epigenetic and transcriptomic human muscles profiles, meta-analyses and OMIC integration, we demonstrated the power of exercise training in shifting the epigenome and transcriptome towards a younger state. We hope that this work will inspire future studies to look deeper at the mechanisms underlying this shift of muscle epigenetic and transcriptomic patterns towards younger profiles.

## Methods

This study was a large-scale investigation of the effect of exercise training on the aging muscle methylome and transcriptome in humans. We used a wide range of bioinformatics and computational techniques (data mining, epigenome-wide association studies, transcriptome-wide association studies, random effects meta-analysis, overrepresentation analysis and multi-contrast enrichment analysis) to analyse and interpret large amounts of OMIC data in human muscle. By exploiting the power of meta-analysis, we overcome many limitations of ‘omics’ research in humans. Specifically, large sample sizes are required to detect changes with small effect sizes, which is the case of age^10,11^ and exercise^32,33,66,67^-related changes in muscle OMIC profiles. All bioinformatics and statistical analyses were performed using the R statistical software.

### Data mining

#### Description of muscle DNA methylation and mRNA expression datasets

First, we gathered all existing DNAm and mRNA expression datasets from our laboratory and our collaborators’, in conjunction with public repositories, to assemble an exhaustive database of DNAm and mRNA expression profiles in muscle (**Fig 1**, Supplementary Tables 1 & 2). We focused exclusively on *microarray* experiments, as they are widely used, scalable (so individual datasets have larger sample sizes), and they measure the same CpGs or transcripts across datasets (so they are straightforward to meta-analyse). We collected the methylomes of 1,251 human samples from 16 datasets, profiled on the Illumina HumanMethylation platform (27K, 450K and EPIC) (Supplementary Table 1), as well as the transcriptomes of 1,926 samples from 21 datasets, profiled on Affimetrix, Illumina, and Agilent platforms (Supplementary Table 2). For robustness, we only included datasets with > 20 samples, with an age SD > 5 years (age-associated changes cannot be detected if age is invariant). Cohorts varied in their age range, health status, ethnicity and potential treatments, so were adjusted for relevant covariates in the statistical analysis to detect age-, CRF-, exercise- and disuse-related changes that are independent of undesirable confounders (see Supplementary Tables 1 & 2 for the list of confounders adjusted in each cohort).

#### Pre-processing

We downloaded the raw IDAT files and pre-processed all DNAm datasets, except for dataset GSE50498 for which we were missing batch information (we used the already pre-processed matrix for this dataset). Details on the pre-processing steps have been previously published^10^, and the preprocessing code is available on <u>Sarah Voisin’s</u> Github account. First, we obtained β-values defined as 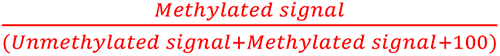. Then, we confirmed the sex of each sample by using the DNAm signal from the sex chromosomes^68^ and removed any sample whose annotated sex did not match predicted sex (3 samples removed across 16 datasets). We used the *ChAMP* pipeline^69^ to preprocess each dataset; we ensured all samples had <10% of probes with detection p-value >0.01, and only excluded probes with missing β-values, with a detection p-value > 0.01, or with a bead count <3 in more than 5% of samples. We removed non-CG probes, SNP-related probes^70^ and probes aligning to multiple locations; for datasets containing males and females, probes located on the sex chromosomes were also removed. Then, a β-mixture quantile normalisation method was applied to adjust for the Type I and Type II probe designs for methylation profiles generated from the HM450 and HMEPIC arrays. To identify technical and biological sources of variation in each individual dataset, singular value decomposition was performed. In all pre-processed datasets, both the plate and the position on the plate were identified as significant technical effects. Thus, all β-values were converted to M-values, and the ComBat function from the *sva* package71 used to adjust directly for these technical artefacts.

We downloaded the already pre-processed mRNA expression datasets, resolved any gene ID ambiguity (e.g. outdated gene names) and averaged the expression of transcripts annotated to the same EntrezID gene.

##### Identifying age-related changes in the muscle methylome and transcriptome

###### EWAS and TWAS meta-analysis of age

To determine whether exercise training can slow down/reverse OMIC aging in human muscle, we first need to know which CpGs and mRNAs change during aging, in which direction, and to what extent. Therefore, we first conducted independent EWAS or TWAS of age in each methylation or transcription dataset. Details on the EWAS pipeline are available elsewhere^10^, and TWAS were conducted in a similar manner. The code for each EWAS and each TWAS is available on <u>Sarah Voisin’s Github account</u>. Briefly, we regressed the DNAm level for each CpG (or the mRNA expression level for each transcript) against age, and adjusted the models for dataset-specific covariates known to influence DNAm or mRNA expression levels (e.g. sex, ethnicity). We then conducted a random-effects meta-analysis to pool the summary statistics at each CpG and mRNA across datasets. Not all CpGs were present in all DNAm datasets, and not all mRNAs were present in all transcriptional datasets, and restricted the analysis to the 595,541 CpG sites present in at least 10 of the 16 DNAm cohorts, and we restricted the analysis to the 16,657 genes present in at least 15 of the 21 datasets. Meta-analysis was carried out using *metafor*72 using “EB” (Empirical Bayes) as the residual heterogeneity estimator, 0.5 as the step length, 10000 iterations, and an accuracy of 1^-8^ in the algorithm that estimates τ^2^. CpGs and mRNAs that showed a meta-analysis false discovery rate (FDR) < 0.00536 were considered age-related and selected for downstream analyses.

###### Over-representation analysis (ORA) of ontologies (molecular pathways, human phenotypes)

To gain insights into the cellular and physiological consequences of aging on the muscle methylome and transcriptome, we tested whether genes belonging to canonical pathways (<u>CP</u> gene set in MSigDB), expression signatures of genetic and chemical perturbations (<u>CGP</u> gene set in MsigDB), gene ontology terms (<u>GO</u> gene set in MsigDB), and human phenotype ontologies (<u>HPO</u> gene set in MsigDB) were over-represented among the age-related CpGs and mRNAs. ORA was performed with the *missmethyl* package^44,45^ for DNAm, using all 595,541 tested CpGs as the background; ORA was performed with the *clusterProfiler* package73 for transcription, using all 16,657 genes as the background. The ORA was restricted to gene sets containing 10-500 genes to limit type I error rate. Gene sets showing an FDR < 0.00536 were considered significantly overrepresented.

###### Confounding by changes in muscle cell type proportions

We used the same ORA technique to estimate whether age-related changes in DNAm signal were potentially confounded by changes in muscle cell type proportions. We created a gene set containing markers genes for muscle cell types identified in a recent single-cell transcriptional study of human muscle^38^ (marker genes for muscle endothelial cells, smooth muscle cells, pericytes, FAP cells, PCV endothelial cells, satellite cells, FBN1 FAP cells, NK cells, myeloid cells, B cells and T cells were from the “Rubenstein_skeletal_muscle” gene set in MSigDB, while marker genes for type I and type II fibers were downloaded directly from the original paper’s supplementary table). Cell types showing an FDR < 0.00536 were considered significantly overrepresented.

###### Integration of aging methylome & transcriptome

We used the same ORA technique to estimate whether there was a significant overlap between age-related changes at DNAm and mRNA expression levels. Age-related differentially methylated genes (DMGs) were used as a gene set in the ORA for transcription, and age-related differentially expressed genes (DEGs) were used as a gene set in the ORA for DNAm.

##### Estimating the effects of CRF, exercise training and muscle disuse on the aging methylome and transcriptome

All analyses described henceforth have been conducted on the age-associated Differentially Methylated Positions (DMPs) and Differentially Expressed genes (DEGs) identified in Step 1.

###### Cardiorespiratory fitness

We focused on the muscle datasets for which information on baseline maximal oxygen uptake (VO_2max_) was available (**Table 1**). We only included datasets with baseline VO_2max_ SD > 5 mL/min/kg (CRF-associated changes cannot be detected if there is no variability in baseline CRF between participant). VO_2max_, measured during a graded exercise test, is considered the gold-standard measurement of CRF.

We applied the same EWAS and TWAS meta-analysis pipeline described previously but regressing DNAm or mRNA expression levels against VO_2max_. We then performed meta-analysis for each aging CpG and transcript across datasets and adjusted for multiple testing. Aging CpGs or mRNAs that were associated with VO_2max_ at FDR < 0.005^36^ were considered significant.

###### Exercise training

For DNAm, we focused on the muscle datasets that came from exercise training studies (**Table 2**). We included all types of exercise training interventions (aerobic training, high-intensity interval training, resistance training) as our aim was to test whether exercise in general could counteract the effect of age on aging OMIC profiles. We applied the same EWAS meta-analysis pipeline described previously but looking at changes in DNAm levels after exercise training (i.e. regressing DNAm levels against Timepoint (PRE/POST training). Then, we performed the meta-analysis for each aging CpG across datasets and adjusted for multiple testing. Aging CpGs whose DNAm levels changed following exercise training at FDR < 0.00536 were considered significant.

For mRNA expression, we extracted summary statistics at age-related mRNAs from a recently published meta-analysis of exercise-induced transcriptional changes^32^. For each transcript, we used the summary statistics from the model selected by the authors (all data came from the meta_analysis_input.RData file uploaded by the authors on <u>their Github page</u>)

###### Muscle disuse

There were no available muscle immobilisation studies that profiled DNAm patterns in human muscle, so we could not estimate the effect of muscle disuse on age-related DNAm patterns.

For mRNA expression, we extracted summary statistics at age-related mRNAs from a published meta-analysis of disuse-induced transcriptional changes^33^. We used summary statistics sent by the authors upon correspondence with them.

##### Transcriptomic integration of age, CRF, exercise and disuse

To contrast the effects of age, CRF, exercise training and muscle disuse, we used multi-contrast enrichment analysis as implemented in the *mitch* package^43^. As recommended in the package, we first created a score to represent the importance of the differential gene expression for each transcript and each contrast:

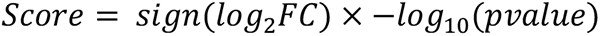

For example, the *FEZ2* gene decreased in expression with age at a magnitude of log_2_FC = -0.12 per year of age at a p-value of 1.24 × 10^−10^. The age score for *FEZ2* was therefore −1 × − log_10_(1.24 × 10^−10^) = −9.9.

We then performed multi-contrast enrichment analysis using the same gene sets as in the ORA analysis described above (canonical pathways, expression signatures of genetic and chemical, gene ontology terms, and human phenotype ontologies), using the default parameters in the mitch_calc() function. Gene sets showing an FDR < 0.005 were considered significant.

##### Visualisation tools

Pairwise correlation plots using Spearman correlations were graphed using the *GGally* package, heatmaps were graphed using the *pheatmap* package after scaling the effect sizes for age, VO_2max_, exercise and/or disuse, and forest plots were graphed using the *metafor* package.

## Data availability

Please refer to Tables 1 & 2 to access all data used in this manuscript. The code producing the Results & Figures for this paper is available on <u>Sarah Voisin’s Github account</u>.

## Acknowledgements

This work was supported by Sarah Voisin’s National Health & Medical Research Council (NHMRC) Early Career Research Fellowship (APP11577321) and by Nir Eynon’s NHMRC Career Development Fellowship (APP1140644). The Gene SMART and LITER studies were both supported by the Collaborative Research Network for Advancing Exercise and Sports Science (201202) scheme awarded to VGC and KJA from the Department of Education and Training, Australia. Mr Nicholas Harvey was supported by a PhD stipend also provided by Bond University CRN-AESS. This research was also supported by infrastructure purchased with Australian Government EIF Super Science Funds as part of the Therapeutic Innovation Australia – Queensland Node project (LRG). Work at Ulster was supported by an Interdisciplinary Award. We also greatly acknowledge Erika Guzman at the ATGC/IHBI/QUT for performing the HMEPIC assays in the LITER study. The EPIK study was supported by the Research Foundation Flanders (FWO G.0898.15).

The authors declare that they have no conflict of interest.

## Author contributions

S.V. performed the bioinformatics and statistical analyses, prepared figures and wrote the manuscript, with contribution from K.S.; S.L., M.J., N.R.H., K.J.A., L.M.H., L.R.G., V.G.C., T.D., J.M.T. collected samples and provided data for the Gene SMART study; M.E.L. provided data for the EpiTrain study; C.W., G.D., R.I. and C.M. provided data for the EXACT study; O.H. and O.A. provided data for the Malmö Prevention and MSAT studies; A.E.H., P.P., K.P. and M.O. provided data for the FTC study; S.B. and M.T. provided data for the EPIK study; C.K.D. provided data for the E-MTAB-11282 study; A.P.S. provided data for the GSE114763 and CAUSE studies; N.E. supervised the work and provided detailed feedback on the analyses and the manuscript. All authors contributed to editing and finalisation of the manuscript.

